# Discovery of long non-coding RNAs in the liver fluke, *Fasciola hepatica*

**DOI:** 10.1101/2023.03.07.531498

**Authors:** Paul McVeigh, Erin McCammick, Emily Robb, Peter Brophy, Russell M Morphew, Nikki J Marks, Aaron G Maule

## Abstract

Long non-coding (lnc)RNAs are a class of eukaryotic RNA that do not code for protein and are linked with transcriptional regulation, amongst a myriad of other functions. Using a custom *in silico* pipeline we have identified 6,436 putative lncRNA transcripts in the liver fluke parasite, *Fasciola hepatica*; none of which are conserved with those previously described from *Schistosoma mansoni. F. hepatica* lncRNAs were distinct from *F. hepatica* mRNAs in transcript length, coding probability, exon/intron composition, expression patterns, and genome distribution. RNA-Seq and digital droplet PCR measurements demonstrated developmentally regulated expression of lncRNAs between intra-mammalian life stages; a similar proportion of lncRNAs (14.2 %) and mRNAs (12.8 %) were differentially expressed (p<0.001), supporting a functional role for lncRNAs in *F. hepatica* life stages. While most lncRNAs (81%) were intergenic, we identified some that overlapped protein coding loci in antisense (13%) or intronic (6%) configurations. We found no unequivocal evidence for correlated developmental expression within positionally correlated lncRNA:mRNA pairs, but global co-expression analysis identified five lncRNA that were inversely co-regulated with 89 mRNAs, including a large number of functionally essential proteases. The presence of micro (mi)RNA binding sites in 3135 lncRNAs indicates the potential for miRNA-based post-transcriptional regulation of lncRNA, and/or their function as competing endogenous (ce)RNAs. This first description of lncRNAs in *F. hepatica* provides an avenue to future functional and comparative genomics studies that will provide a new perspective on a poorly understood aspect of parasite biology.

## 1. INTRODUCTION

*Fasciola* spp. liver fluke are helminth parasites that impact the health and productivity of farm animals, leading to considerable costs for international agricultural economies (Spithill, 1999; Gray, 2008; NADIS, 2016). *Fasciola* also infects humans and is recognized as a zoonotic Neglected Tropical Disease (NTD) pathogen (Hotez et al., 2008). Fluke pathogenicity is compounded by the prevalence of resistance alleles for most flukicidal drugs (Kelley et al., 2016) and increased prevalence driven by predictions of warmer, wetter, weather patterns associated with the climate crisis (Alba et al., 2021).

Efforts to develop new treatments for liver fluke infections have been supported by the development of systems-level experimental resources over the past decade, including genome, transcriptome, proteome and functional genomics methodologies for *F. hepatica* (McVeigh et al., 2018). This increased volume and resolution of ‘omics data has yielded insights into the non-coding (nc)RNAs of *Fasciola* genomes, where (mi)RNA complements are beginning to be clarified (Xu et al., 2012; Fontenla et al., 2015; Fromm et al., 2015; Guo and Guo, 2019; Ovchinnikov et al., 2020; Ricafrente et al., 2020; Herron, 2021; Hu et al., 2021), including a subset that appears to be secreted within extracellular vesicles (EVs).

Transfer RNA fragments are the only other ncRNA to have been identified from *F. hepatica* transcriptomes (Hu et al., 2021), but eukaryotic transcriptomes contain a plethora of ncRNA biotypes. One of the most widely studied over recent years has been the long non-coding (lnc)RNAs, arbitrarily defined as transcripts longer than 200nt that do not code for protein. Interest in this family has largely been driven by perceived importance in human tumour biology (Cossu et al., 2019; Lecerf et al., 2019; Zhao et al., 2019), however a burgeoning research field also describes these molecules in non-human animal systems, as well as plants, yeast and prokaryotes (Brown et al., 1992; Bernstein et al., 1993; Clemson et al., 1996; Reeves et al., 2007; Houseley et al., 2008; Swiezewski et al., 2009). Amongst lncRNA functions are roles in transcription, translation, protein localization, cellular integrity, the cell cycle, apoptosis, and stem cell pluripotency (for review see (Ma et al., 2013)). These functions were identified by RNA interference (RNAi) experiments, focusing on cytosolic lncRNAs in human systems (Goyal et al., 2017; Salehi et al., 2017), and increasingly, CRISPR-based perturbation of lncRNA expression in human cell lines (Qi et al., 2019), and mice (Butler et al., 2019). This volume of functional data means that mammalian experimental systems remain our touchstone for understanding lncRNA biology. Functional genomics methods have been less widely applied to invertebrates, but RNAi has yielded functional data in arthropods (Etebari et al., 2016; Yang et al., 2016) while CRISPR has yielded lncRNA functional insights in *C. elegans* (Akay et al., 2019; Wei et al., 2019). While no data on lncRNA functions are available from phylum Platyhelminthes, the first lncRNA sequence datasets have now been published from parasitic flatworms, including *Schistosoma mansoni* and *S. japonicum* (Vasconcelos et al., 2017; Liao et al., 2018; Oliveira et al., 2018; Kim et al., 2020; Maciel et al., 2020), *Echinococcus granulosus* (Zhang et al., 2020) and *Macrostomum lignano* (Azlan et al., 2020). These molecules are now being recognised as potential therapeutic targets for new anti-parasitic drugs (Silveira et al., 2022) This paper provides the first description of long non-coding (lnc)RNAs in *F. hepatica*. These data provide important insights into potential functions for lncRNAs and will form the basis for future functional genomics studies. Our hope is that these data will trigger new avenues towards studying, and new perspectives on, liver fluke biology.

## 2 METHODS

### 2.1 Transcriptome assemblies

Workflow was as described in Figure 1. We began by assembling a non-redundant transcriptome from available *F. hepatica* RNA-Seq datasets. These included previously-published (Cwiklinski et al., 2018), non-stranded, paired-end Illumina sequencing read sets from *F. hepatica* metacercariae (met; *n=*3), newly excysted juveniles (NEJ) (1h (nej1h; *n=*2), 3h (nej3h; *n=*2), 24h (nej24h; *n=*2) *in vitro*), *in vivo* liver stage parasites recovered from rats at 21 days post oral exposure (juv21d; *n=*1), adult parasites (Ad; *n=*1), and eggs (egg; *n=*1), with fastq files obtained from the European Nucleotide Archive (ENA) via project PRJEB6904). We supplemented these with stranded paired-end Illumina sequencing datasets for rat-derived juv21d worms (juv21d; *n=3*), and 21 day *in vitro* maintained worms (invitro21d; *n*=3). These six libraries are available from ENA project PRJEB49655, samples ERS9656160-ERS9656165, generated as described in (Robb et al., 2022). The *F. gigantica* 18h NEJ RNA-Seq dataset was generated as described in (Davey et al., 2022), available from DDBJ/EMBL/GenBank accession GJHP01000000 and can be interrogated *via* https://sequenceserver.ibers.aber.ac.uk/.

**Figure 1.**
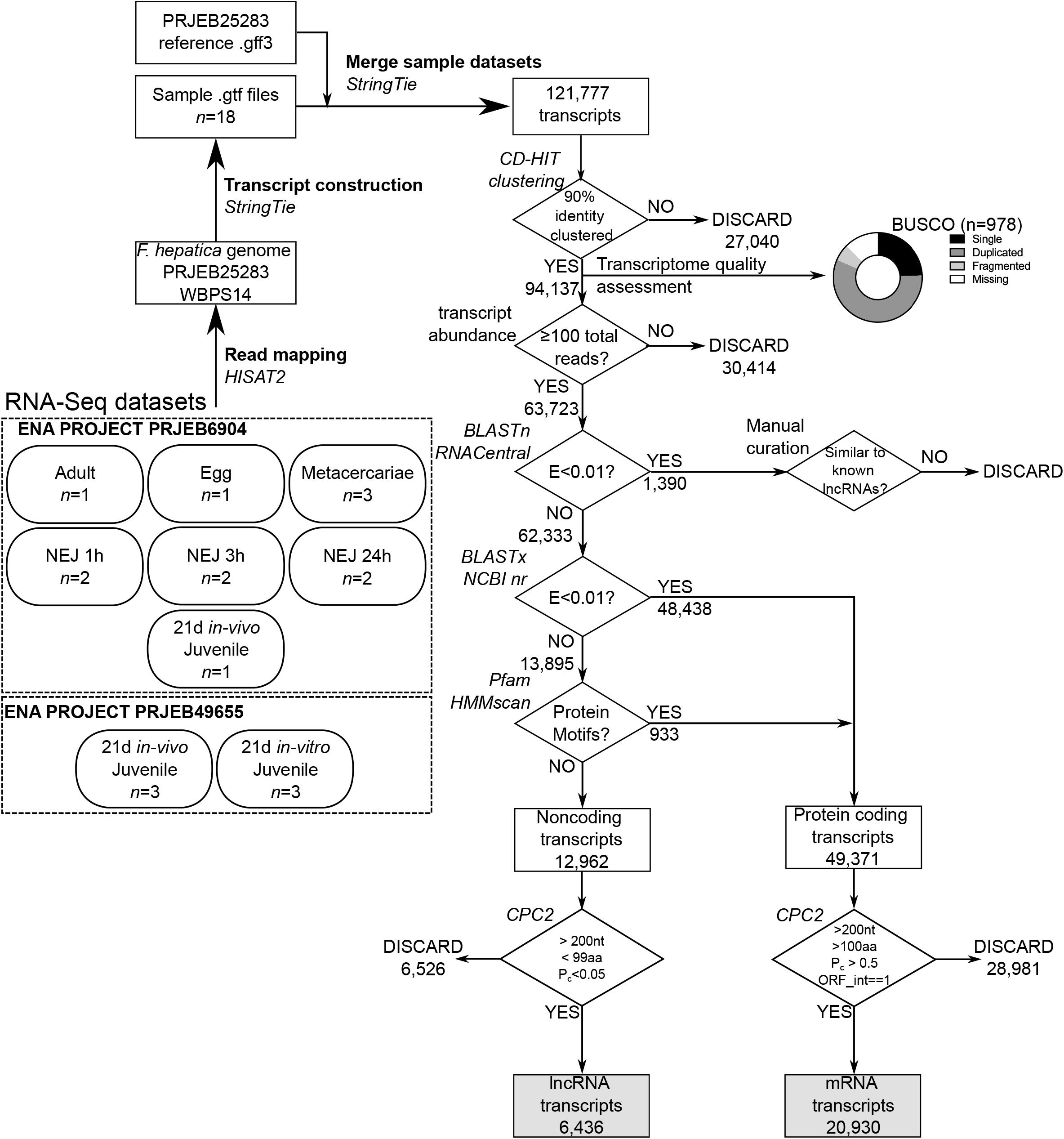
Long non-coding RNA discovery pipeline. Flowchart indicates processing and analysis of RNA-Seq Illumina short read datasets into lncRNA data. Numbers of sequences passing or failing each filter stage of the pipeline are indicated, software tools are named in italics.

Fastq files were mapped against the *F. hepatica* genome (WormBase ParaSite PRJEB25283, database version WBPS14) using HISAT2, with a merged transcript assembly generated through StringTie (as described in (Pertea et al., 2016); using default parameters throughout). This produced 120,927 transcripts of _≧_200 nt length (the default length cutoff within StringTie), which were passed into our lncRNA annotation pipeline (Figure 1). A separate stranded transcriptome was also produced by mapping the three (Robb et al., 2022) juv21d datasets against the *F. hepatica* genome as described above, and employing the -*-fr* option in StringTie to denote a stranded library in “fr-firststrand” format, matching the dUTP-based synthesis method of the stranded library synthesis kit that we used (Robb et al., 2022).

### 2.2 Identification of *F. hepatica* lncRNAs

After removal of duplicate transcripts (Figure 1), and application of an expression cutoff of _≧_100 reads per transcript, contaminating non-coding sequences (ribosomal and transfer RNAs) were removed by comparison with the RNAcentral dataset (The, 2019). Forty sequences possessing weak similarity with known lncRNAs were replaced back into the dataset. The remaining 63,805 sequences were filtered for similarity with proteins using BLASTx, performed locally using DIAMOND BLAST (Buchfink et al., 2015) against the ncbi nr protein sequence database. Transcripts scoring E<0.01 (44,713 sequences) were filtered into a ‘coding sequence’ bin. Remaining sequences (19,092) were screened for protein motifs using Pfam hmmscan, performed locally, resulting in removal of an additional 455 transcripts with motif hits above the default inclusion threshold. The remaining 18,637 presumed non-coding sequences were all analysed with Coding Potential Calculator (CPC2 (Kang et al., 2017)) in a process of manual cross-checking. This final step ensured that our lncRNAs all had ‘lncRNA-like’ characteristics (>200nt length, encoding _≧_100 aa peptide, defined by CPC2 as “noncoding”, with a coding probability of <0.05). Our final dataset contained 7,497 lncRNA transcripts from 6,321 genomic loci. Figure 1 also describes a parallel pipeline for identification of coding RNAs (presumed “mRNAs”), all of which were finally assessed by CPC2 as ‘coding’, with coding probability _≧_0.95, with transcript length _≧_200nt, and encoded peptides _≧_100 aa), yielding 21,697 mRNA transcripts from 9,046 loci.

### 2.3 Differential expression analyses

To identify transcripts with statistically significant differential expression (DE) between sequential life cycle stages, we used pairwise exact tests within the EdgeR package (Robinson et al., 2010; McCarthy et al., 2012). Only life stages supported by at least 2 biological replicates were used for these tests (pairwise tests performed: met vs nej1h; neje1h vs nej3h; nej3h vs nej24h; nej24h vs juv21d; juv21d vs met). Raw count data, recovered from StringTie using the prepDE.py script described in the StringTie manual, were piped into EdgeR for analysis. EdgeR output files were parsed using custom Python scripts for list comprehension, allowing extraction of fold change data supported by p_≧_0.001.

### 2.4 Digital droplet PCR

To confirm RNA-Seq transcript expression data, we performed digital droplet (dd)PCR on six randomly selected lncRNAs across three life stages. Total RNA was extracted using Trizol (ThermoFisher) from one adult *F. hepatica* (*n*=3), and from the same juv21d (*n=3*) and invitro21d (*n*=3) RNA samples used for transcriptome sequencing as described in section 2.1. Each cDNA was generated from 500ng total RNA using the High Capacity RNA-to-cDNA kit (Thermo Fisher). Amplification for ddPCR used the EvaGreen Supermix (BioRad), including 200 nM of each primer (Supporting file 9). Data analysis used the Quantasoft package from BioRad.

### 2.5 Genome localisation and expression correlation

For identification of antisense and intronic lncRNAs we used the bedtools package (v2.28; (Quinlan and Hall, 2010). After generating bed files for transcripts, exons, introns and intergenic regions from our gtf file, we used bedtools *intersect* function to identify overlapping lncRNA and mRNA transcript loci, and sorted these by strand orientation (based on our stranded juv21d datasets) to identify antisense lncRNA:mRNA pairs. Bedtools *intersect* again allowed us to identify intronic lncRNAs by comparing lncRNA exons with mRNA introns, and again, these were sorted by strand orientation. Intergenic lncRNAs were identified as those that were not amongst our lists of antisense or intronic lncRNAs. We used the bedtools *closest* algorithm to identify the single closest upstream and downstream mRNA locus to each intergenic lncRNA locus.

Before examining expression correlation, we parsed all antisense, intronic and intergenic lncRNA:mRNA pairs to identify those pairs in which both members were DE, using a Python script for list comprehension. We then used the ‘CORREL’ function in MS Excel to calculate the correlation coefficient for lncRNA vs mRNA across TPM expression data from met, nej1h, nej3h, nej24h and juv21d libraries.

### 2.6 miRNA response element identification

To examine whether any of our lncRNAs contained binding sites for miRNAs (miRNA response elements; MRE), we employed a consensus prediction method as described by (Gillan et al., 2017). Working with the 150 miRNAs previously described in *F. hepatica* (Xu et al., 2012; Fontenla et al., 2015; Fromm et al., 2015; Ovchinnikov et al., 2020; Herron, 2021), we employed three miRNA target prediction tools to identify binding matches between all *F. hepatica* miRNAs and our lncRNAs; we retained only those lncRNA:miRNA pairs that were identified by all three algorithms. We used local instances of miRANDA (Enright et al., 2003), RNAhybrid (Rehmsmeier et al., 2004) and PITA (Kertesz et al., 2007), accepting hits fulfilling the thresholds as used by (Gillan et al., 2017): miRanda, total score >145, energy < -10; RNAhybrid, p<0.1, energy < -22; PITA, _≧_G < -10. LncRNA:miRNA pairs identified by all three algorithms were extracted using a custom Python script, and only these consensus pairs were accepted.

## 3. RESULTS

### 3.1 Dataset Summary

Mapping of the >1.7 bn reads (81.7% overall alignment rate; Supplementary Data Sheet 1) associated with 15 RNA-Seq libraries yielded 121,777 transcripts, which clustered at 90 % identity into 94,137 non-redundant transcripts (Figure 1). Using BUSCO (Seppey et al., 2019), we compared the quality of our assembly with the *F. hepatica* transcript assemblies available through WormBase Parasite WBPS14. Our dataset contained more complete BUSCOs (81.7%; PRJNA179522 = 40.4%, PRJEB25283 = 74.1%), and had fewer missing BUSCOs (12.5%; PRJNA179522 = 31.3%, PRJEB25283 = 17.9%), than previous assemblies (Figure 1).

At this point, we filtered the transcriptome to include only sequences represented by at least 100 reads, leaving a total of 63,723 supported transcripts (Supplementary Data Sheet 2).

From this dataset, we removed 1,390 sequences with similarity to other classes of ncRNA (ribosomal RNA, transfer RNA etc) listed in the RNACentral dataset. We then extracted protein coding transcripts (containing BLAST or Pfam identities, and a complete ORF) into a counterpart dataset containing 20,930 “mRNA” transcripts (Supplementary Data Sheet 3), which we used to compare and examine the transcriptional and syntenic relationships between lncRNA and mRNA loci. Having removed mRNAs, we extracted our putative lncRNAs (6,436 transcripts measuring _≧_200 nt and lacking significant protein coding capacity; Supplementary Data Sheets 4-6).

### 3.2 lncRNA and mRNA datasets are quantitatively and qualitatively distinct

The pipeline described in Figure 1 identified 6,436 transcripts fitting the definition of lncRNAs, and 20,930 transcripts with protein coding capacity, which we label here as mRNAs. These datasets were distinct on all measures examined (Figure 2) – lncRNA transcripts were shorter overall (median: lncRNA=875 nt, mRNA=3133 nt; Figure 2A), encoded shorter open reading frames (median: lncRNA=49 aa, mRNA=397 aa; Figure 2B), displayed lower protein coding probability (median: lncRNA P=0.025, mRNA P=1.000; Figure 2C), and were expressed more scarcely (median: lncRNAs 0.122 TPM, mRNA=0.628 TPM; Figure 2D) than mRNA transcripts. Loci coding for lncRNAs were smaller than mRNA loci (median: lncRNA=2010 bp, mRNA=29,261 bp; Figure 2E), with longer exons (median: lncRNA=319 bp, mRNA=169 bp; Figure 2F) and shorter introns (median: lncRNA=322 bp, mRNA=1412 bp; Figure 2G). LncRNA loci incorporated fewer exons (median=2 exons; Figure 2H), compared to mRNA loci (median=7 exons; Figure 2I).

**Figure 2.**
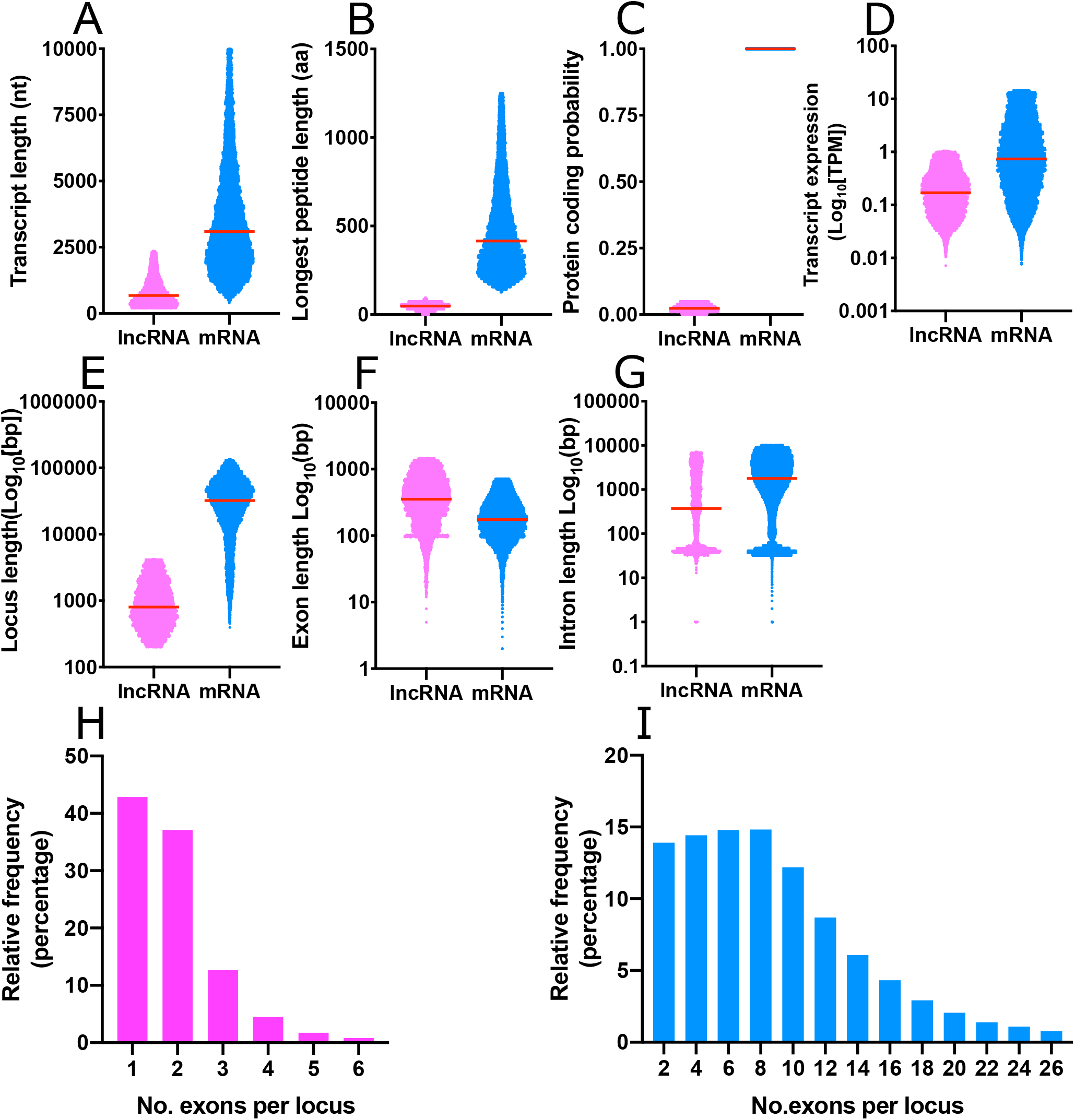
Statistical summary of mRNA and long noncoding (lnc)RNA annotations from *Fasciola hepatica*. lncRNAs and mRNAs are qualitatively and quantitatively distinct across all measures examined: (A) Transcript length; (B) Open reading frame length; (C) Probability of protein coding capacity; (D) Transcript expression/abundance; (E) Genomic locus length; (F) Exon length; (G) Intron length; and (H, I) Number of exons per locus. Each scatter graph (A-G) is composed of individual datapoints, with dataset median illustrated by a horizontal red bar. In all graphs, lncRNAs are magenta, mRNA are blue.

### 3.3 *hepatica* lncRNAs are dissimilar to lncRNAs from non-*Fasciola* species

To identify lncRNA orthologues in the closely related species, *F. gigantica*, we used BLASTn to compare our 6,436 *F. hepatica* lncRNA transcripts with an *F. gigantica* NEJ transcriptome of 16,551 transcripts This identified 204 potential lncRNA orthologues (Supplementary Data Sheet 7, 8). These putative lncRNAs were (median±SD) 551±267 nt length, and incorporated 144±84 nt ORFs. All showed >80% sequence identity with an *F. hepatica* lncRNA, and all were designated “non-coding” by Coding Potential Assessment Tool (CPAT).

We also compared *F. hepatica* lncRNAs with published lncRNAs from other flatworms for where those sequences were openly available. These included the >10,000 lncRNA transcripts reported from *S. mansoni* (Vasconcelos et al., 2017; Maciel et al., 2019). CD-HIT identified no evidence for lncRNA sequence similarity between *F. hepatica* and *S. mansoni*, while BLASTn identified a single, low-scoring orthologous pair, matching the *F. hepatica* lncRNA STRG.25709.1 with SmLINC02629-IBu from *S. mansoni* (bitscore 56.5, E-value 4e^-07^). Likewise, BLASTn comparison with *M. lignano* lncRNAs (Azlan et al., 2020) found no similarity, while comparisons with the RNAcentral non-coding RNA database found no similarity with the >200,000 mammalian lncRNAs within that dataset.

### 3.4 lncRNA transcripts are differentially expressed during development

Differential expression (DE) analyses employed only life stages represented by at least two biological replicate libraries (met, nej1h, nej3h, nej24h, juv21d, invitro21d). Figure 3 and Supplementary Data Sheet 9 show that all of these libraries bore stage-specific lncRNAs (met=4; nej1h=4; nej3h=3; nej24h=1; juv21d=63; invitro21d=13), and mRNAs (met=42; nej1h=35; nej3h=45; nej24h=30; juv21d=225; invitro21d=97). Using edgeR’s exact test algorithm, pairwise comparisons were performed between developmentally sequential life stages, defining DE transcripts as those with a statistically significant difference (p_≧_0.001) in at least one of these comparisons (Figure 3A; Supplementary Data Sheet 10). A total of 911 DE lncRNA transcripts (Figure 3B; 14.2% of all lncRNAs), and 2,673 DE mRNA transcripts (Figure 3C; 12.8% of all mRNAs) were identified. DE transcripts were found in every pairwise comparison (Figure 3D). To confirm these data, six randomly selected lncRNAs were tested for expression across adult, juv21d and invitro21d life stages using ddPCR; in every case the ddPCR expression pattern correlated well with the NGS expression pattern (Supplementary Data Sheet 11).

**Figure 3.**
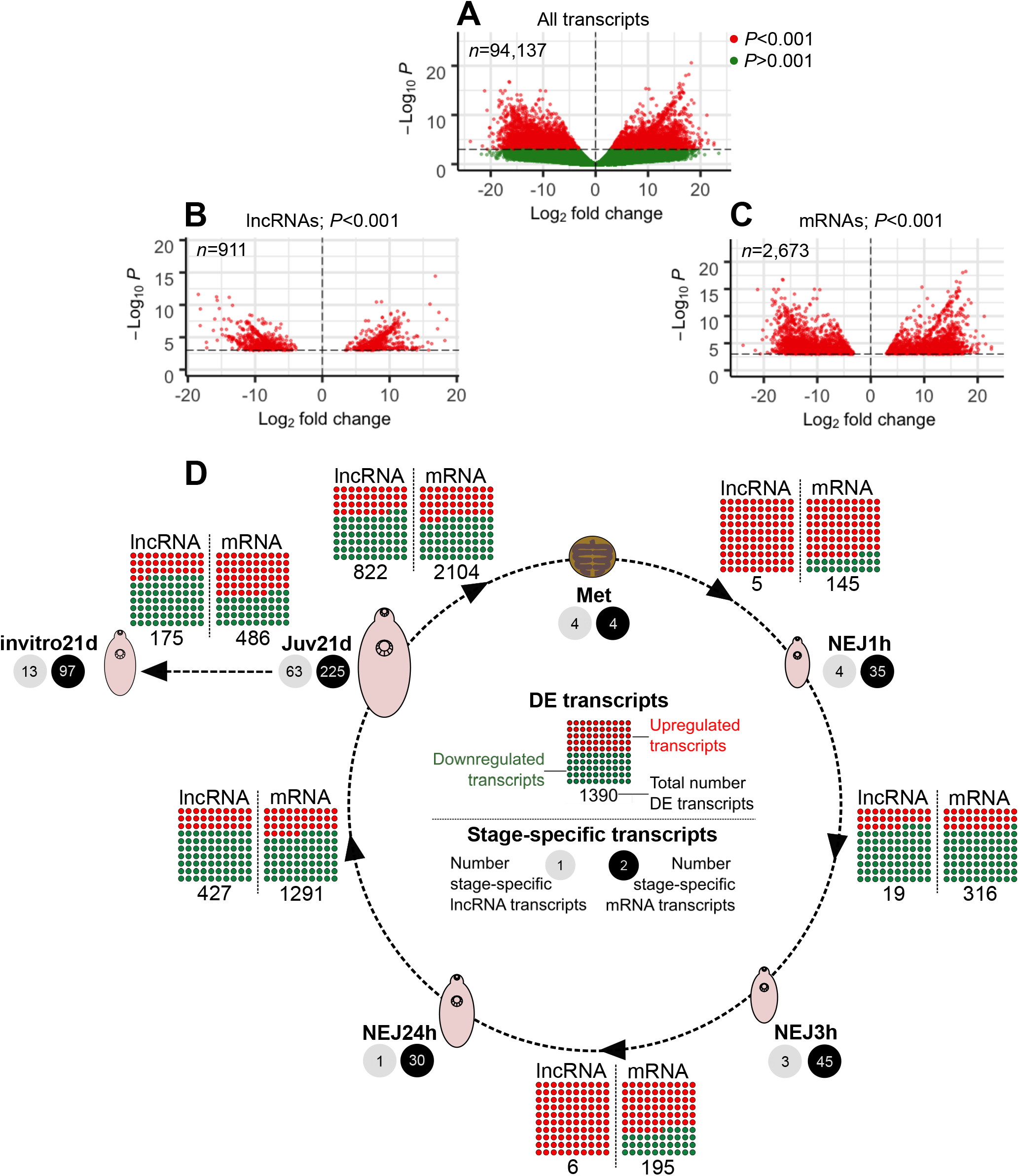
Developmental regulation of long non-coding (lnc)RNAs in intra-mammalian stages of Fasciola hepatica. Statistically significant differential expression (DE) was determined for lncRNA and mRNA transcripts during developmental transitions, showing that F. hepatica lncRNAs are dynamically-regulated during intra-mammalian development. A-C. Volcano plots showing distribution of transcript fold change vs P value across all comparisons. A, all transcripts, with P-value cutoff as indicated on y-axis; B, Fold change distribution of DE lncRNAs; C, Fold change distribution of DE mRNAs. D, Summary data from DE analysis of transcripts in metacercariae (Met), newly excysted juvenile (NEJ) maintained in vitro for 1h post excystment (NEJ1h), 3h post excystment (NEJ3h) or 24h post excystment (NEJ24h), and ex vivo parasites recovered from rat livers at 21 days post infection (Juv21d). Numbers of stage-specific lncRNAs and mRNAs at each life stage are indicated, as are the numbers of DE lncRNA and mRNA transcripts, and the proportion of each upregulated or downregulated, between each developmental transition.

### 3.5 Positional relationships and expression correlations between lncRNAs and mRNAs

Based on genomic location and directionality relative to protein-coding loci, we identified three lncRNA types in *F. hepatica*: (i) antisense lncRNAs: expressed from a lncRNA exon on the opposite DNA strand to a protein-coding locus, and overlapping a protein-coding exon by at least 1bp; (ii) intronic lncRNAs: expressed from lncRNA exons in either orientation that reside within an intron of a protein coding locus and do not overlap an exon; (iii) intergenic lncRNAs: expressed from distinct loci that do not intersect with any protein coding locus. To delineate our lncRNAs into these categories, we used the bedtools software package, and stranded juv21d libraries (as described in Methods). Our first approach was to search for sense:antisense overlaps between lncRNA and mRNA transcripts using the bedtools *intersect* algorithm, which yielded 536 non-redundant lncRNA:mRNA transcript pairs, overlapping in opposite transcriptional orientations. The same approach identified 534 similarly oriented mRNA:mRNA pairs, and 17 lncRNA:lncRNA pairs (Supplementary Data Sheet 12)

Parsing the data to identify antisense pairs in which both members showed statistically significant differential expression (DE) identified 14 DE lncRNA:mRNA pairs and five DE mRNA:mRNA pairs. There were no lncRNA:lncRNA pairs in which both members were DE (Supplementary Data Sheet 12). For all pairs we calculated the Pearson correlation coefficient (CC) of lncRNA vs mRNA expression (TPM) across all life stages (Supplementary Data Sheet 12). Accepting >0.99 and <-0.99 as cut-offs identified just 53 pairs (10 % of the total number of pairs) of “expression correlated” lncRNA/mRNA transcript pairs.

After removing antisense transcripts from the dataset, we identified 30 intronic lncRNA:mRNA pairs (median CC = 0.407; Supplementary Data Sheet 13). No pairs consisted of both members as DE, and none passed the +/-0.9 CC cut-off. Finally, we identified 3795 intergenic lncRNAs who’s closest transcript was an mRNA (Supplementary Data Sheet 14). In 29 pairs both members were DE, and only three pairs passed the +/-0.9 CC cut-off.

To explore *trans* interactions in more detail, we employed a weighted correlation network analysis approach. Here, we sought evidence for co-expression between all lncRNAs and all mRNAs across all libraries with multiple biological replicates. CEMiTool identified a single co-expression module, containing 89 mRNAs and five lncRNAs (Supplementary Data Sheet 15). The mean TPM of the lncRNAs and mRNAs within this module were inversely correlated across longitudinally sampled libraries (Figure 4), suggesting that they may be co-regulated.

**Figure 4.**
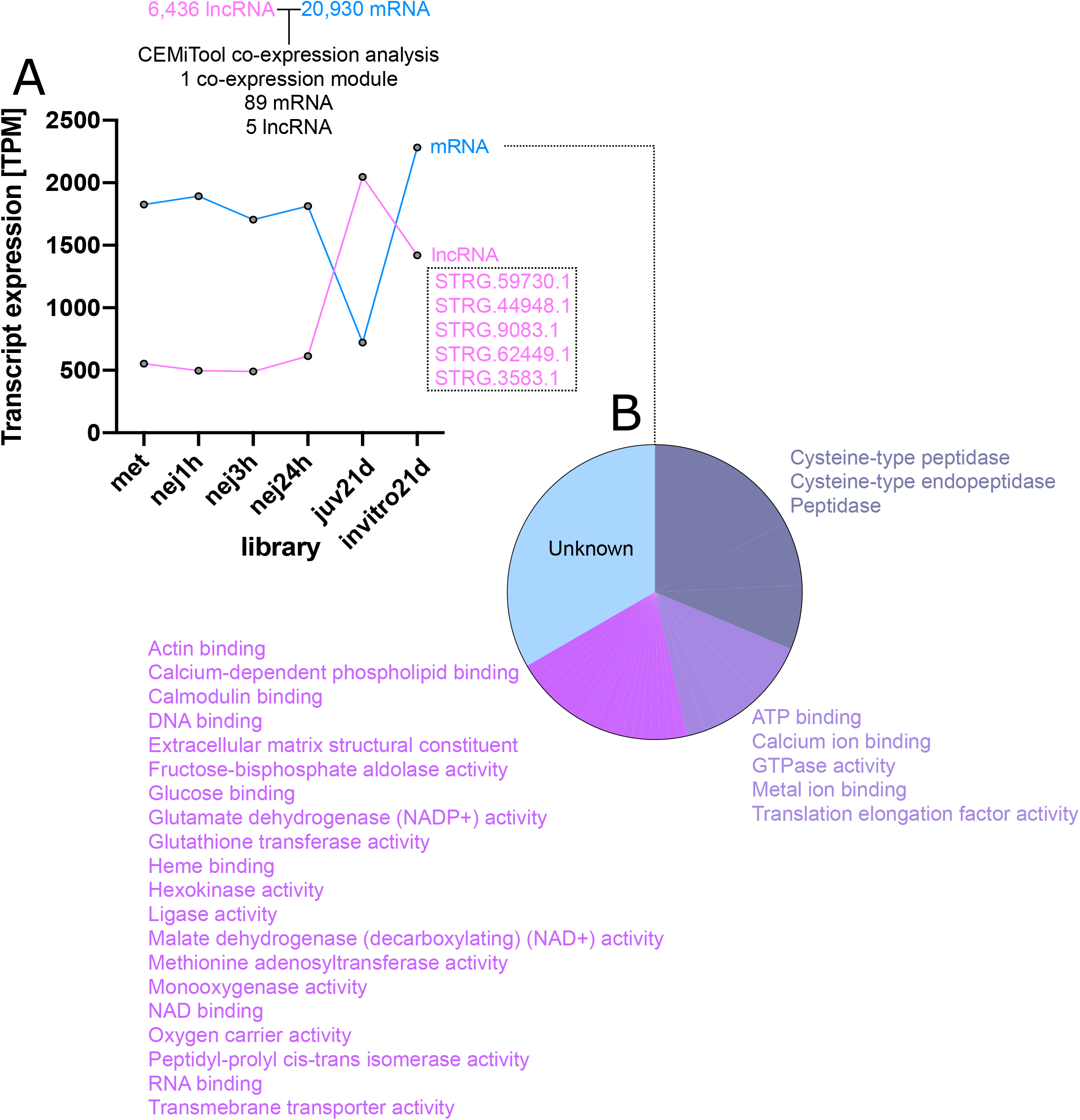
Co-expression of lncRNAs and mRNAs. (A) Global co-expression analysis of all lncRNAs and mRNAs in our dataset. CEMiTool identified one co-expression module in which five lncRNAs and 89 mRNAs show inversely correlated expression across life stage RNA-Seq libraries. (B) The 89 mRNAs comprise mostly cysteine proteases in addition to a range of other sequences as indicated. The remaining one third of matching mRNAs code for unknown sequence types that lacked BLAST homology and identifiable sequence motifs.

### 3.6 *F. hepatica* lncRNAs contain miRNA binding sites

LncRNAs contain binding sites for miRNAs, implying either that they serve as “sponges” for “soaking up” individual miRNAs, or that their expression is regulated by miRNA binding and transcriptional destruction. Previous reports have identified at least 150 miRNAs in *F. hepatica* (Xu et al., 2012; Fontenla et al., 2015; Fromm et al., 2015; Ovchinnikov et al., 2020; Herron, 2021). Using three miRNA binding prediction tools (miRANDA, PITA, RNAhybrid) we analysed these miRNAs in the context of binding to our lncRNAs, producing a consensus set of 4104 lncRNA:miRNA pairs that were present in outputs from all three tools (we rejected any matches that were not predicted by all three tools). Supplementary Data Sheet 16 shows that these matches incorporated all 150 miRNAs, and 2618 lncRNAs. Individual lncRNAs contained binding sites for up to eight distinct miRNAs, with each miRNA predicted to bind up to 234 lncRNAs. Output data from PITA illustrated the number of individual binding sites per individual lncRNA:miRNA pair: while 95 % of lncRNAs contained ten or fewer binding sites, 5% displayed up to 246. Figure 5 shows an excerpt of this lncRNA:miRNA network, focusing on the lncRNAs matching eight or more miRNAs. Within this network, STRG.61441.1 was the major hub lncRNA, binding to eight miRNAs. Fhe-pubnovelmiR-7 was the major hub miRNA, binding to 234 lncRNAs.

**Figure 5.**
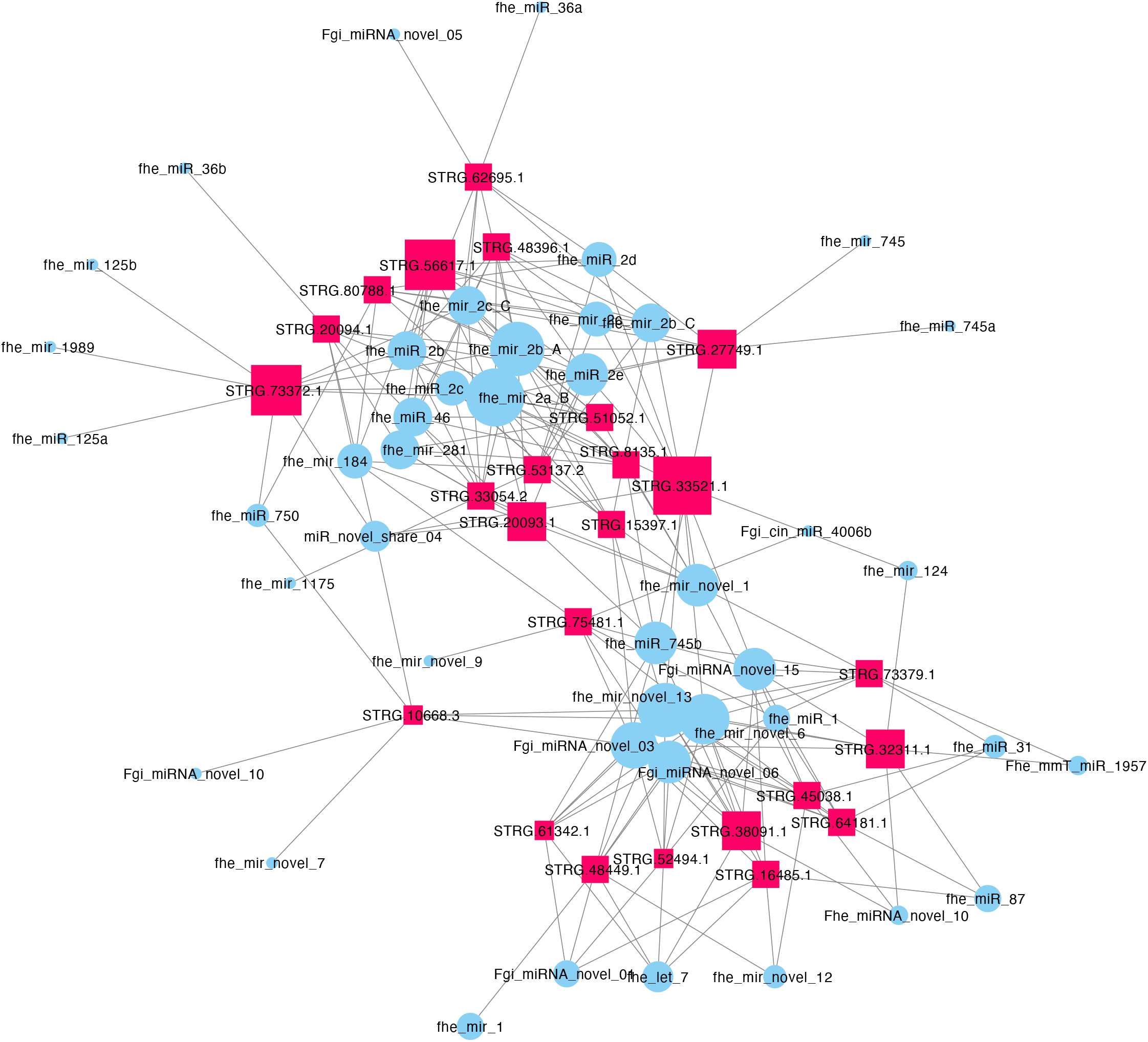
Network representation of computationally predicted binding interactions between Fasciola hepatica long noncoding (lnc)RNAs and micro (mi)RNAs. This network represents an excerpt of the lncRNA:miRNA network detailed in Supporting File 12, showing only the lncRNAs that matched eight or more miRNAs. LncRNA nodes are represented by red squares, miRNA nodes by blue circles. Node size is proportional to the number of connections with other nodes. Connections between nodes are represented by grey edges.

## 4 DISCUSSION

Fasciolosis, caused by *F. hepatica* and *F. gigantica*, is an important veterinary and zoonotic disease which requires improved diagnostic and control approaches if it is to be sustainably combatted in the long term. A key approach to identification of new control targets and diagnostic biomarkers is through increased understanding of fundamental parasite biology and host-parasite interactions. In this work, we have addressed this need by focusing on long non-coding RNAs in *F. hepatica* for the first time. This study has performed the first classification of lncRNAs in *F. hepatica*, linking these to predicted roles in parasite development, stem cell biology, and co-regulation of miRNAs and mRNAs. This work represents an essential precursor to functional understanding of liver fluke lncRNA biology.

Unlike more mature systems, *F. hepatica* does not yet have a unified description of the protein-coding transcriptome that would have enabled a simple subtraction-based approach to lncRNA discovery. Disparities can be seen in the varying gene complements presented by currently available studies (Cwiklinski et al., 2015; McNulty et al., 2017; Cwiklinski et al., 2018). This meant that we needed to generate a consensus transcriptome before filtering lncRNAs from mRNAs within those datasets. Our consensus transcriptome combined 18 publicly available and newly generated libraries, which together encompassed 94,137 non-redundant transcripts. This represented by far the largest transcriptome yet described for *F. hepatica*, with previous datasets describing 22,676 (Cwiklinski et al., 2015; Cwiklinski et al., 2018), 16,806 (WormBase ParaSite release WBPS14), or 14,462 (McNulty et al., 2017) transcripts, although it is unclear whether those previously published datasets retained non-coding sequences. BUSCO metrics confirmed that our transcriptome assembly was the most complete yet published for *Fasciola*, giving us confidence in proceeding to lncRNA discovery. This transcriptome was then filtered to include only sequences represented by at least 100 reads (an arbitrary selection which aimed to balance well-supported transcript models while maintaining adequate detection of rare transcripts), followed by removal of irrelevant classes of ncRNA, and protein coding transcripts. Our final pool of putative lncRNAs represented 6,436 transcripts measuring _≧_200 nt and lacking significant protein coding capacity. A separate pool of protein-coding transcripts contained 20,930 mRNAs.

To identify lncRNA transcripts that we may have mis-annotated, or those from published datasets that may have been erroneously annotated as protein-coding, we used BLAST to compare all lncRNAs with previously published *F. hepatica* transcript annotations. BLASTx did not identify any identical matches between our lncRNAs and previously predicted proteins. BLASTn analyses identified 37 lncRNAs matching 20 sequences previously annotated as mRNAs. However, we found that these matching sequences had either no identifiable protein sequence characteristics and/or were classed as hypothetical proteins. We therefore retained these in our lncRNA dataset as putative lncRNAs.

Previous literature from other species showed that lncRNAs were distinct from mRNAs across a wide range of measures (Etebari et al., 2016; Feng et al., 2018; Wang et al., 2018; Zhou et al., 2018). We confirmed that these findings also applied to our dataset, showing that lncRNAs were shorter than mRNAs in gene locus length, overall transcript length and open reading frame length. LncRNA loci also tended to display fewer, longer exons and shorter introns than mRNA loci. LncRNAs were also expressed more scarcely than mRNA transcripts. Inter-species comparisons of nucleic acid sequences can provide insights into evolution, with sequence conservation potentially indicating functional conservation. There are currently limited opportunities to make this comparison for lncRNAs with other flatworms, with *S. mansoni, S. japonicum, E. granulosus* and *M. lignano* the only other flatworm lncRNA datasets currently available (Vasconcelos et al., 2017; Liao et al., 2018; Oliveira et al., 2018; Azlan et al., 2020; Zhang et al., 2020). In comparing *F. hepatica* lncRNAs with these datasets we found an almost complete lack of lncRNA sequence conservation. A previous study did report lncRNA sequence conservation between *S. mansoni, S. japonicum* and *S. haematobium*, albeit with lower similarity than that seen in mRNA comparisons (Liao et al., 2018). To our knowledge, ours is the first inter-genus comparison of lncRNAs in phylum Platyhelminthes. This absence of primary sequence similarity is not surprising and suggests that, like in other genera, the rapid evolution of lncRNAs invalidates primary sequence similarity as a means of identifying lncRNA homologs (Diederichs, 2014; Johnsson et al., 2014; Hezroni et al., 2015). Similarly, BLASTn comparisons with the RNAcentral non-coding RNA database found no similarity between our lncRNAs and the >200,000 mammalian lncRNAs within that dataset. Rather than primary sequence similarity, lncRNAs are thought to be conserved between species according to spatiotemporal and syntenic locus expression (Jarroux et al., 2017), criteria which will require more mature genome assemblies before they can be properly analysed in flatworms.

Differential expression (DE) analyses employed only life stages represented by at least two biological replicate libraries (met, nej1h, nej3h, nej24h, juv21d, invitro21d). All these libraries bore stage-specific lncRNAs and mRNAs, with 911 DE lncRNA transcripts and 2,673 DE mRNA transcripts identified in pairwise comparisons between sequential lifestages. DE transcripts were found in all pairwise comparisons, with the number of DE transcripts roughly proportional to the time-course length between stages – the fewest between met:nej1h (5 lncRNA, 145 mRNA), and the most between juv21d:met (822 lncRNA, 2104 mRNA).

Comparison of juv21d with invitro21d samples (respectively, flukes recovered from rat livers 21 days after infection, or maintained *in vitro* for 21 days (McCusker et al., 2016; Robb et al., 2022), demonstrated the presence of 13 lncRNAs found uniquely in invitro21d samples, with 63 found uniquely in juv21d worms. Given the key developmental differences between these groups (Robb et al., 2022), these lncRNAs have potential importance for fluke development, and represent priority targets for functional genomics experiments. The strikingly similar proportions of DE lncRNAs and mRNAs (respectively, 14.2% and 12.8%), shows that lncRNAs are at least as transcriptionally dynamic as coding RNAs, and supports the hypothesis that they have essential and stage-specific functions during the *F. hepatica* intra-mammalian life cycle.

While our focus was not to reannotate the mRNA transcriptome of *F. hepatica*, our lncRNA annotation method did also generate protein-coding mRNAs and associated DE data.

Briefly, the most highly regulated mRNA transcripts (_≧_16 fold) in our dataset closely reflect previous observations (Cwiklinski et al., 2018), and include sequences within GO processes describing cell adhesion, cell division, glycolysis, metal ion homeostasis, muscle function, proteolysis, protein synthesis and modification, RNA transcriptional control and signal transduction. The purpose of these data was to identify and explore potential interactions between lncRNAs and mRNAs. Various classification schemes exist for lncRNAs, but one of the most commonly used groups sequences based on their genomic location and directionality relative to protein-coding loci (Jarroux et al., 2017). According to this classification scheme, we used a subtractive approach to identify antisense, intronic and intergenic lncRNA types in *F. hepatica*. These classifications are important because they can inform potential interactions with protein coding genes, for example *cis* natural antisense transcripts (NATs) and intronic lncRNAs can affect the expression of corresponding sense transcripts from their “host” gene, while *trans*-antisense and intergenic lncRNAs can impact expression of distant genomic loci. Our searches yielded 536 antisense overlapping lncRNA and mRNA transcript pairs (where the mRNA was considered “sense”). These data also identified 534 complementary mRNA:mRNA pairs, and 17 complementary lncRNA:lncRNA pairs. These data show that lncRNA and/or mRNA loci may overlap on the genome, to produce pairs of antisense/complementary-oriented transcripts. Given evidence that lncRNAs can interact transcriptionally in *cis* or *trans* fashion with mRNAs, we looked for evidence of transcriptional interaction between antisense transcripts. For example, data from human cell lines show that antisense lncRNAs can regulate the expression of the mRNAs with which they overlap (Balbin et al., 2015; Jadaliha et al., 2018). We reasoned that correlated expression might be most clearly identified in cases where both members of an antisense pair were DE. Parsing the data to identify pairs in which this was the case identified 14 DE lncRNA:mRNA pairs. We also found five DE mRNA:mRNA pairs, but no lncRNA:lncRNA pairs in which both members were DE. In case these criteria were prohibitively restrictive, we also calculated the Pearson correlation coefficient (CC) of lncRNA vs mRNA expression (TPM) for all 536 pairs across all life stages. Accepting >0.99 and <-0.99 as cut-offs identified just 53 pairs of DE lncRNA/mRNA transcript pairs that were highly correlated. These similar data suggested that correlated expression between *cis* oriented antisense transcripts might not be a widespread phenomenon, but we cannot rule out the possibility that some antisense lncRNA regulation of *cis* mRNA transcripts may occur. Functional genomics experiments will be necessary to test hypotheses around linkages between individual transcripts, and separate interacting transcripts from physically overlapping transcripts that do not interact (Karlin et al., 2002; Veeramachaneni et al., 2004; Kim et al., 2009; Behura and Severson, 2013). As well as informing studies on evolution of the *F. hepatica* genome, these data have important practical implications in avoiding off-target effects in RNAi or CRISPR/Cas9 experimental design. After removing these antisense transcripts from the dataset, we used similar methods to identify 30 intronic lncRNA:mRNA pairs (i.e. lncRNAs located within an mRNA intron in either orientation). No pair consisted of both members as DE, and none passed the +/-0.9 CC cut-off. Finally, we identified 3795 intergenic lncRNAs who’s closest transcript was an mRNA (Supporting file 12). In 29 pairs both members were DE, and only three pairs passed the +/-0.9 CC cut-off. These findings differ from those from other systems, where lncRNAs have been shown to affect transcription of antisense overlapping and neighbouring protein-coding genes (Balbin et al., 2015; Liu et al., 2017; Jadaliha et al., 2018). This prompted us to explore *trans* lncRNA interactions in more detail, where lncRNAs may regulate distant mRNA loci through epigenetic interaction (Ransohoff et al., 2018). We approached this using weighted correlation network analysis, via the CEMiTool R package, focusing broadly on co-expression between all lncRNAs and all mRNAs across all libraries in our datasets. This identified a single co-expression module, containing 89 mRNAs and five lncRNAs, which showed inversely correlated expression suggesting interaction and/or co-regulation.

Interestingly, 47 % of mRNA transcripts within the module were associated with peptidase activity, including Cathepsin L and B, and legumain (Figure 4). This is unsurprising given the preponderance of proteases amongst the most highly regulated *F. hepatica* transcripts (Cwiklinski et al., 2015; Cwiklinski et al., 2018); these data represent the first link between lncRNAs and potential co-regulation of protease expression. Future experiments should cement this link by silencing one or more of the lncRNAs and assaying for fluctuation in linked protease transcripts.

One of the most widespread hypotheses for lncRNA function is the competing endogenous (ce)RNA hypothesis, which holds that lncRNAs contain miRNA binding sites (known as miRNA Response Elements, or MREs), enabling them to act as ‘sponges’ for miRNAs. This ‘sponging’ phenomenon is thought to promote competition for miRNA binding with cognate mRNA targets, enabling fine control of miRNA regulation of mRNA target transcripts (Thomson and Dinger, 2016; Ulitsky, 2018). Alternatively, the presence of miRNA binding sites may simply indicate that lncRNAs are transcriptionally regulated by miRNA binding in the same manner as mRNAs (Yamamura et al., 2018). Since both possibilities could yield important insights into liver fluke lncRNA biology, we used in silico tools to explore the presence of miRNA binding sites on lncRNAs.

One hundred and fifty-nine miRNAs have been previously reported in *F. hepatica* (Xu et al., 2012; Fontenla et al., 2015; Fromm et al., 2015; Ovchinnikov et al., 2020; Herron, 2021). We generated in silico binding predictions between these miRNAs and our lncRNAs, producing a consensus set of 4104 lncRNA:miRNA pairs. These matches incorporated all 150 miRNAs, and 2618 lncRNAs. Individual lncRNAs contained binding sites for up to eight distinct miRNAs, with each miRNA predicted to bind up to 234 lncRNAs. Output data from PITA illustrated the number of individual binding sites per individual lncRNA:miRNA pair: while 95 % of lncRNAs contained ten or fewer binding sites, the remaining 5% displayed up to 246. Network analysis identified STRG.61441.1 as the major hub lncRNA, binding to eight miRNAs. Fhe-pubnovelmiR-7 was the major hub miRNA, binding to 234 lncRNAs. These data support the possibility of miRNA-lncRNA interactions, that might manifest as either traditional miRNA-driven post-transcriptional regulation of lncRNA expression, or a ceRNA function for the described miRNAs. Experimental evidence will be required to test these hypotheses further.

We have profiled the first set of lncRNAs in the liver fluke, *F. hepatica*. These non-coding RNAs were expressed across multiple intra-mammalian developmental stages, showing dynamic regulation between life-stages that suggests life-stage specific functions. *In silico* analyses supported important roles for lncRNAs in transcriptional regulation including: (i) an inverse correlation of lncRNA expression with mRNAs, suggesting co-regulation of these sequences, and; (ii) the widespread location of miRNA binding sites on lncRNAs, suggesting miRNA regulation of lncRNA, or vice versa. These data represent a steppingstone towards an understanding of non-coding RNA biology in *F. hepatica*, an area which remains poorly understood across eukaryotes but could expose new therapeutic and diagnostic options for parasite infections globally.

## Supporting information

Supplementary Data Sheet 1

Supplementary Data Sheet 2

Supplementary Data Sheet 3

Supplementary Data Sheet 4

Supplementary Data Sheet 5

Supplementary Data Sheet 6

Supplementary Data Sheet 7

Supplementary Data Sheet 8

Supplementary Data Sheet 9

Supplementary Data Sheet 10

Supplementary Data Sheet 11

Supplementary Data Sheet 12

Supplementary Data Sheet 13

Supplementary Data Sheet 14

Supplementary Data Sheet 15

Supplementary Data Sheet 16

