## Supplementary Data Sheet 1 for "Discovery of long non-coding RNAs in the liver fluke, *Fasciola hepatica*"

```
# SUPPLEMENTARY DATA SHEET 1. HISAT2 ALIGNMENT STATISTICS OF ILLUMINA RNA-SEQ READS DURING MAPPING TO THE FASCIOLA HEPATICA GENOME (PRJEB25283, WBPS14)


37008210 reads; of these:
  37008210 (100.00%) were paired; of these:
    5824380 (15.74%) aligned concordantly 0 times
    28767490 (77.73%) aligned concordantly exactly 1 time
    2416340 (6.53%) aligned concordantly >1 times
    ----
    5824380 pairs aligned concordantly 0 times; of these:
      394120 (6.77%) aligned discordantly 1 time
    ----
    5430260 pairs aligned 0 times concordantly or discordantly; of these:
      10860520 mates make up the pairs; of these:
        6811084 (62.71%) aligned 0 times
        3568089 (32.85%) aligned exactly 1 time
        481347 (4.43%) aligned >1 times
90.80% overall alignment rate


88178696 reads; of these:
  88178696 (100.00%) were paired; of these:
    19088333 (21.65%) aligned concordantly 0 times
    62767953 (71.18%) aligned concordantly exactly 1 time
    6322410 (7.17%) aligned concordantly >1 times
    ----
    19088333 pairs aligned concordantly 0 times; of these:
      637098 (3.34%) aligned discordantly 1 time
    ----
    18451235 pairs aligned 0 times concordantly or discordantly; of these:
      36902470 mates make up the pairs; of these:
        27381548 (74.20%) aligned 0 times
        8289920 (22.46%) aligned exactly 1 time
        1231002 (3.34%) aligned >1 times
84.47% overall alignment rate


66495596 reads; of these:
  66495596 (100.00%) were paired; of these:
    15804958 (23.77%) aligned concordantly 0 times
    45721937 (68.76%) aligned concordantly exactly 1 time
    4968701 (7.47%) aligned concordantly >1 times
    ----
    15804958 pairs aligned concordantly 0 times; of these:
      758960 (4.80%) aligned discordantly 1 time
    ----
    15045998 pairs aligned 0 times concordantly or discordantly; of these:
      30091996 mates make up the pairs; of these:
        24120797 (80.16%) aligned 0 times
        5130382 (17.05%) aligned exactly 1 time
        840817 (2.79%) aligned >1 times
81.86% overall alignment rate


66662513 reads; of these:
  66662513 (100.00%) were paired; of these:
    12063790 (18.10%) aligned concordantly 0 times
    49673950 (74.52%) aligned concordantly exactly 1 time
    4924773 (7.39%) aligned concordantly >1 times
    ----
    12063790 pairs aligned concordantly 0 times; of these:
      642272 (5.32%) aligned discordantly 1 time
    ----
    11421518 pairs aligned 0 times concordantly or discordantly; of these:
      22843036 mates make up the pairs; of these:
        16059838 (70.31%) aligned 0 times
        5841050 (25.57%) aligned exactly 1 time
        942148 (4.12%) aligned >1 times
87.95% overall alignment rate


100376452 reads; of these:
  100376452 (100.00%) were paired; of these:
    22181196 (22.10%) aligned concordantly 0 times
    70526638 (70.26%) aligned concordantly exactly 1 time
    7668618 (7.64%) aligned concordantly >1 times
    ----
    22181196 pairs aligned concordantly 0 times; of these:
      679835 (3.06%) aligned discordantly 1 time
    ----
    21501361 pairs aligned 0 times concordantly or discordantly; of these:
      43002722 mates make up the pairs; of these:
        32392772 (75.33%) aligned 0 times
        9279345 (21.58%) aligned exactly 1 time
        1330605 (3.09%) aligned >1 times
83.86% overall alignment rate


83454099 reads; of these:
  83454099 (100.00%) were paired; of these:
    15306507 (18.34%) aligned concordantly 0 times
    62223929 (74.56%) aligned concordantly exactly 1 time
    5923663 (7.10%) aligned concordantly >1 times
    ----
    15306507 pairs aligned concordantly 0 times; of these:
      1276019 (8.34%) aligned discordantly 1 time
    ----
    14030488 pairs aligned 0 times concordantly or discordantly; of these:
      28060976 mates make up the pairs; of these:
        19293590 (68.76%) aligned 0 times
        7560582 (26.94%) aligned exactly 1 time
        1206804 (4.30%) aligned >1 times
88.44% overall alignment rate


63496634 reads; of these:
  63496634 (100.00%) were paired; of these:
    13438696 (21.16%) aligned concordantly 0 times
    43925920 (69.18%) aligned concordantly exactly 1 time
    6132018 (9.66%) aligned concordantly >1 times
    ----
    13438696 pairs aligned concordantly 0 times; of these:
      682934 (5.08%) aligned discordantly 1 time
    ----
    12755762 pairs aligned 0 times concordantly or discordantly; of these:
      25511524 mates make up the pairs; of these:
        20017103 (78.46%) aligned 0 times
        4770963 (18.70%) aligned exactly 1 time
        723458 (2.84%) aligned >1 times
84.24% overall alignment rate


83491240 reads; of these:
  83491240 (100.00%) were paired; of these:
    12930553 (15.49%) aligned concordantly 0 times
    63048373 (75.51%) aligned concordantly exactly 1 time
    7512314 (9.00%) aligned concordantly >1 times
    ----
    12930553 pairs aligned concordantly 0 times; of these:
      792040 (6.13%) aligned discordantly 1 time
    ----
    12138513 pairs aligned 0 times concordantly or discordantly; of these:
      24277026 mates make up the pairs; of these:
        15764300 (64.94%) aligned 0 times
        7351469 (30.28%) aligned exactly 1 time
        1161257 (4.78%) aligned >1 times
90.56% overall alignment rate


45746545 reads; of these:
  45746545 (100.00%) were paired; of these:
    9707089 (21.22%) aligned concordantly 0 times
    32850942 (71.81%) aligned concordantly exactly 1 time
    3188514 (6.97%) aligned concordantly >1 times
    ----
    9707089 pairs aligned concordantly 0 times; of these:
      324670 (3.34%) aligned discordantly 1 time
    ----
    9382419 pairs aligned 0 times concordantly or discordantly; of these:
      18764838 mates make up the pairs; of these:
        14054261 (74.90%) aligned 0 times
        4129898 (22.01%) aligned exactly 1 time
        580679 (3.09%) aligned >1 times
84.64% overall alignment rate


45540680 reads; of these:
  45540680 (100.00%) were paired; of these:
    10265697 (22.54%) aligned concordantly 0 times
    32127883 (70.55%) aligned concordantly exactly 1 time
    3147100 (6.91%) aligned concordantly >1 times
    ----
    10265697 pairs aligned concordantly 0 times; of these:
      313277 (3.05%) aligned discordantly 1 time
    ----
    9952420 pairs aligned 0 times concordantly or discordantly; of these:
      19904840 mates make up the pairs; of these:
        14680781 (73.75%) aligned 0 times
        4602315 (23.12%) aligned exactly 1 time
        621744 (3.12%) aligned >1 times
83.88% overall alignment rate


72247393 reads; of these:
  72247393 (100.00%) were paired; of these:
    12245612 (16.95%) aligned concordantly 0 times
    54574139 (75.54%) aligned concordantly exactly 1 time
    5427642 (7.51%) aligned concordantly >1 times
    ----
    12245612 pairs aligned concordantly 0 times; of these:
      835588 (6.82%) aligned discordantly 1 time
    ----
    11410024 pairs aligned 0 times concordantly or discordantly; of these:
      22820048 mates make up the pairs; of these:
        15473606 (67.81%) aligned 0 times
        6347665 (27.82%) aligned exactly 1 time
        998777 (4.38%) aligned >1 times
89.29% overall alignment rate


44738981 reads; of these:
  44738981 (100.00%) were paired; of these:
    14387024 (32.16%) aligned concordantly 0 times
    13280387 (29.68%) aligned concordantly exactly 1 time
    17071570 (38.16%) aligned concordantly >1 times
    ----
    14387024 pairs aligned concordantly 0 times; of these:
      95336 (0.66%) aligned discordantly 1 time
    ----
    14291688 pairs aligned 0 times concordantly or discordantly; of these:
      28583376 mates make up the pairs; of these:
        26268885 (91.90%) aligned 0 times
        1107055 (3.87%) aligned exactly 1 time
        1207436 (4.22%) aligned >1 times
70.64% overall alignment rate


50620773 reads; of these:
  50620773 (100.00%) were paired; of these:
    9324490 (18.42%) aligned concordantly 0 times
    32237254 (63.68%) aligned concordantly exactly 1 time
    9059029 (17.90%) aligned concordantly >1 times
    ----
    9324490 pairs aligned concordantly 0 times; of these:
      673537 (7.22%) aligned discordantly 1 time
    ----
    8650953 pairs aligned 0 times concordantly or discordantly; of these:
      17301906 mates make up the pairs; of these:
        14430545 (83.40%) aligned 0 times
        2106318 (12.17%) aligned exactly 1 time
        765043 (4.42%) aligned >1 times
85.75% overall alignment rate


59478285 reads; of these:
  59478285 (100.00%) were paired; of these:
    10978259 (18.46%) aligned concordantly 0 times
    38409140 (64.58%) aligned concordantly exactly 1 time
    10090886 (16.97%) aligned concordantly >1 times
    ----
    10978259 pairs aligned concordantly 0 times; of these:
      914005 (8.33%) aligned discordantly 1 time
    ----
    10064254 pairs aligned 0 times concordantly or discordantly; of these:
      20128508 mates make up the pairs; of these:
        17568067 (87.28%) aligned 0 times
        1875106 (9.32%) aligned exactly 1 time
        685335 (3.40%) aligned >1 times
85.23% overall alignment rate


33907880 reads; of these:
  33907880 (100.00%) were paired; of these:
    11605120 (34.23%) aligned concordantly 0 times
    10599773 (31.26%) aligned concordantly exactly 1 time
    11702987 (34.51%) aligned concordantly >1 times
    ----
    11605120 pairs aligned concordantly 0 times; of these:
      105046 (0.91%) aligned discordantly 1 time
    ----
    11500074 pairs aligned 0 times concordantly or discordantly; of these:
      23000148 mates make up the pairs; of these:
        21126015 (91.85%) aligned 0 times
        892555 (3.88%) aligned exactly 1 time
        981578 (4.27%) aligned >1 times
68.85% overall alignment rate


59637024 reads; of these:
  59637024 (100.00%) were paired; of these:
    11433673 (19.17%) aligned concordantly 0 times
    38597211 (64.72%) aligned concordantly exactly 1 time
    9606140 (16.11%) aligned concordantly >1 times
    ----
    11433673 pairs aligned concordantly 0 times; of these:
      710246 (6.21%) aligned discordantly 1 time
    ----
    10723427 pairs aligned 0 times concordantly or discordantly; of these:
      21446854 mates make up the pairs; of these:
        18155366 (84.65%) aligned 0 times
        2528113 (11.79%) aligned exactly 1 time
        763375 (3.56%) aligned >1 times
84.78% overall alignment rate


61562095 reads; of these:
  61562095 (100.00%) were paired; of these:
    11806245 (19.18%) aligned concordantly 0 times
    39879200 (64.78%) aligned concordantly exactly 1 time
    9876650 (16.04%) aligned concordantly >1 times
    ----
    11806245 pairs aligned concordantly 0 times; of these:
      922252 (7.81%) aligned discordantly 1 time
    ----
    10883993 pairs aligned 0 times concordantly or discordantly; of these:
      21767986 mates make up the pairs; of these:
        18955776 (87.08%) aligned 0 times
        2099755 (9.65%) aligned exactly 1 time
        712455 (3.27%) aligned >1 times
84.60% overall alignment rate


37417336 reads; of these:
  37417336 (100.00%) were paired; of these:
    12619986 (33.73%) aligned concordantly 0 times
    12136040 (32.43%) aligned concordantly exactly 1 time
    12661310 (33.84%) aligned concordantly >1 times
    ----
    12619986 pairs aligned concordantly 0 times; of these:
      135712 (1.08%) aligned discordantly 1 time
    ----
    12484274 pairs aligned 0 times concordantly or discordantly; of these:
      24968548 mates make up the pairs; of these:
        22980280 (92.04%) aligned 0 times
        937091 (3.75%) aligned exactly 1 time
        1051177 (4.21%) aligned >1 times
69.29% overall alignment rate


51097023 reads; of these:
  51097023 (100.00%) were paired; of these:
    9126319 (17.86%) aligned concordantly 0 times
    33264787 (65.10%) aligned concordantly exactly 1 time
    8705917 (17.04%) aligned concordantly >1 times
    ----
    9126319 pairs aligned concordantly 0 times; of these:
      550609 (6.03%) aligned discordantly 1 time
    ----
    8575710 pairs aligned 0 times concordantly or discordantly; of these:
      17151420 mates make up the pairs; of these:
        14458187 (84.30%) aligned 0 times
        2050084 (11.95%) aligned exactly 1 time
        643149 (3.75%) aligned >1 times
85.85% overall alignment rate


61978384 reads; of these:
  61978384 (100.00%) were paired; of these:
    11105522 (17.92%) aligned concordantly 0 times
    40533815 (65.40%) aligned concordantly exactly 1 time
    10339047 (16.68%) aligned concordantly >1 times
    ----
    11105522 pairs aligned concordantly 0 times; of these:
      713138 (6.42%) aligned discordantly 1 time
    ----
    10392384 pairs aligned 0 times concordantly or discordantly; of these:
      20784768 mates make up the pairs; of these:
        18336738 (88.22%) aligned 0 times
        1849555 (8.90%) aligned exactly 1 time
        598475 (2.88%) aligned >1 times
85.21% overall alignment rate


58960697 reads; of these:
  58960697 (100.00%) were paired; of these:
    21307336 (36.14%) aligned concordantly 0 times
    16198350 (27.47%) aligned concordantly exactly 1 time
    21455011 (36.39%) aligned concordantly >1 times
    ----
    21307336 pairs aligned concordantly 0 times; of these:
      141566 (0.66%) aligned discordantly 1 time
    ----
    21165770 pairs aligned 0 times concordantly or discordantly; of these:
      42331540 mates make up the pairs; of these:
        39250243 (92.72%) aligned 0 times
        1366050 (3.23%) aligned exactly 1 time
        1715247 (4.05%) aligned >1 times
66.71% overall alignment rate


[bam_sort_core] merging from 6 files and 1 in-memory blocks...
58586906 reads; of these:
  58586906 (100.00%) were paired; of these:
    13135775 (22.42%) aligned concordantly 0 times
    40905467 (69.82%) aligned concordantly exactly 1 time
    4545664 (7.76%) aligned concordantly >1 times
    ----
    13135775 pairs aligned concordantly 0 times; of these:
      782491 (5.96%) aligned discordantly 1 time
    ----
    12353284 pairs aligned 0 times concordantly or discordantly; of these:
      24706568 mates make up the pairs; of these:
        21308428 (86.25%) aligned 0 times
        2917152 (11.81%) aligned exactly 1 time
        480988 (1.95%) aligned >1 times
81.81% overall alignment rate


55213746 reads; of these:
  55213746 (100.00%) were paired; of these:
    10906243 (19.75%) aligned concordantly 0 times
    40538711 (73.42%) aligned concordantly exactly 1 time
    3768792 (6.83%) aligned concordantly >1 times
    ----
    10906243 pairs aligned concordantly 0 times; of these:
      757052 (6.94%) aligned discordantly 1 time
    ----
    10149191 pairs aligned 0 times concordantly or discordantly; of these:
      20298382 mates make up the pairs; of these:
        17591860 (86.67%) aligned 0 times
        2351745 (11.59%) aligned exactly 1 time
        354777 (1.75%) aligned >1 times
84.07% overall alignment rate


51707525 reads; of these:
  51707525 (100.00%) were paired; of these:
    10292768 (19.91%) aligned concordantly 0 times
    37716113 (72.94%) aligned concordantly exactly 1 time
    3698644 (7.15%) aligned concordantly >1 times
    ----
    10292768 pairs aligned concordantly 0 times; of these:
      756939 (7.35%) aligned discordantly 1 time
    ----
    9535829 pairs aligned 0 times concordantly or discordantly; of these:
      19071658 mates make up the pairs; of these:
        17157004 (89.96%) aligned 0 times
        1669269 (8.75%) aligned exactly 1 time
        245385 (1.29%) aligned >1 times
83.41% overall alignment rate


41730900 reads; of these:
  41730900 (100.00%) were paired; of these:
    15025966 (36.01%) aligned concordantly 0 times
    11493387 (27.54%) aligned concordantly exactly 1 time
    15211547 (36.45%) aligned concordantly >1 times
    ----
    15025966 pairs aligned concordantly 0 times; of these:
      82673 (0.55%) aligned discordantly 1 time
    ----
    14943293 pairs aligned 0 times concordantly or discordantly; of these:
      29886586 mates make up the pairs; of these:
        27645330 (92.50%) aligned 0 times
        1015204 (3.40%) aligned exactly 1 time
        1226052 (4.10%) aligned >1 times
66.88% overall alignment rate


49328505 reads; of these:
  49328505 (100.00%) were paired; of these:
    11133104 (22.57%) aligned concordantly 0 times
    33982399 (68.89%) aligned concordantly exactly 1 time
    4213002 (8.54%) aligned concordantly >1 times
    ----
    11133104 pairs aligned concordantly 0 times; of these:
      531497 (4.77%) aligned discordantly 1 time
    ----
    10601607 pairs aligned 0 times concordantly or discordantly; of these:
      21203214 mates make up the pairs; of these:
        18729602 (88.33%) aligned 0 times
        2110636 (9.95%) aligned exactly 1 time
        362976 (1.71%) aligned >1 times
81.02% overall alignment rate


55993924 reads; of these:
  55993924 (100.00%) were paired; of these:
    12515871 (22.35%) aligned concordantly 0 times
    38602816 (68.94%) aligned concordantly exactly 1 time
    4875237 (8.71%) aligned concordantly >1 times
    ----
    12515871 pairs aligned concordantly 0 times; of these:
      696025 (5.56%) aligned discordantly 1 time
    ----
    11819846 pairs aligned 0 times concordantly or discordantly; of these:
      23639692 mates make up the pairs; of these:
        21479185 (90.86%) aligned 0 times
        1840451 (7.79%) aligned exactly 1 time
        320056 (1.35%) aligned >1 times
80.82% overall alignment rate


43048853 reads; of these:
  43048853 (100.00%) were paired; of these:
    14784292 (34.34%) aligned concordantly 0 times
    25530175 (59.31%) aligned concordantly exactly 1 time
    2734386 (6.35%) aligned concordantly >1 times
    ----
    14784292 pairs aligned concordantly 0 times; of these:
      246856 (1.67%) aligned discordantly 1 time
    ----
    14537436 pairs aligned 0 times concordantly or discordantly; of these:
      29074872 mates make up the pairs; of these:
        24824375 (85.38%) aligned 0 times
        3671656 (12.63%) aligned exactly 1 time
        578841 (1.99%) aligned >1 times
71.17% overall alignment rate


42792167 reads; of these:
  42792167 (100.00%) were paired; of these:
    15126640 (35.35%) aligned concordantly 0 times
    24960535 (58.33%) aligned concordantly exactly 1 time
    2704992 (6.32%) aligned concordantly >1 times
    ----
    15126640 pairs aligned concordantly 0 times; of these:
      241512 (1.60%) aligned discordantly 1 time
    ----
    14885128 pairs aligned 0 times concordantly or discordantly; of these:
      29770256 mates make up the pairs; of these:
        25094416 (84.29%) aligned 0 times
        3868944 (13.00%) aligned exactly 1 time
        806896 (2.71%) aligned >1 times
70.68% overall alignment rate


38595509 reads; of these:
  38595509 (100.00%) were paired; of these:
    6946407 (18.00%) aligned concordantly 0 times
    28433043 (73.67%) aligned concordantly exactly 1 time
    3216059 (8.33%) aligned concordantly >1 times
    ----
    6946407 pairs aligned concordantly 0 times; of these:
      379535 (5.46%) aligned discordantly 1 time
    ----
    6566872 pairs aligned 0 times concordantly or discordantly; of these:
      13133744 mates make up the pairs; of these:
        9194503 (70.01%) aligned 0 times
        3408483 (25.95%) aligned exactly 1 time
        530758 (4.04%) aligned >1 times
88.09% overall alignment rate


38304766 reads; of these:
  38304766 (100.00%) were paired; of these:
    7356769 (19.21%) aligned concordantly 0 times
    27821998 (72.63%) aligned concordantly exactly 1 time
    3125999 (8.16%) aligned concordantly >1 times
    ----
    7356769 pairs aligned concordantly 0 times; of these:
      373160 (5.07%) aligned discordantly 1 time
    ----
    6983609 pairs aligned 0 times concordantly or discordantly; of these:
      13967218 mates make up the pairs; of these:
        9634977 (68.98%) aligned 0 times
        3759935 (26.92%) aligned exactly 1 time
        572306 (4.10%) aligned >1 times
87.42% overall alignment rate
```
