## Supplementary Data Sheet 11 for "Discovery of long non-coding RNAs in the liver fluke, *Fasciola hepatica*"

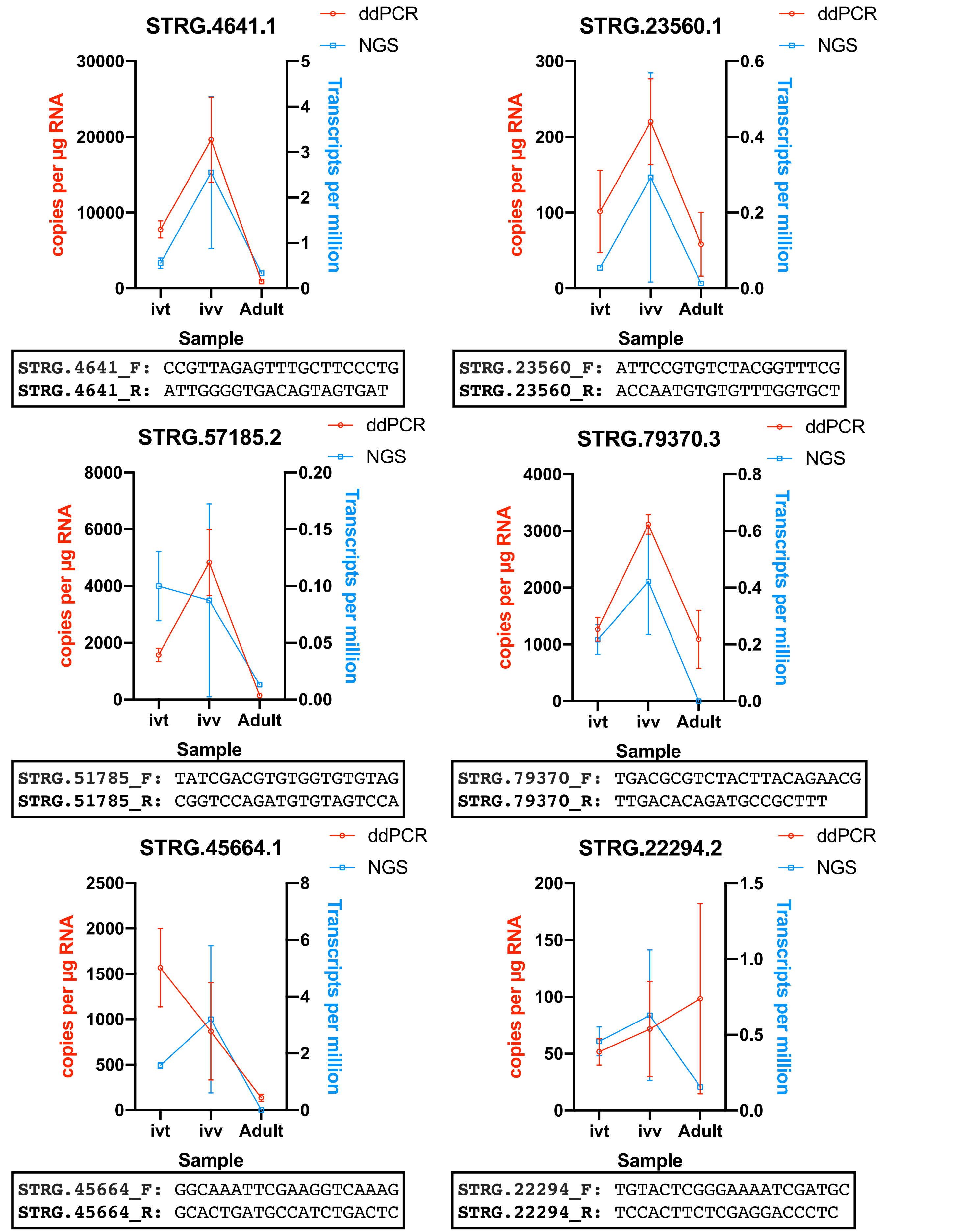


**Supporting File 9. Comparison of expression level of six long non-coding (lnc)RNAs measured with next generation sequencing (NGS) (blue), and digital droplet (dd)PCR (red)**. Each lncRNA was measured across three life-stage libraries: Adult *Fasciola hepatica*, 21 day in vitro juvenile *F. hepatica* (ivt) and 21 day in vivo juvenile *F. hepatica* (ivv). Each datapoint represents the mean±SEM of at least three biological replicates. Under each graph, the primer sets used for ddPCR amplification are indicated.
